# Jovian enables direct inference of germline haplotypes from short reads via sequence-to-sequence modeling

**DOI:** 10.1101/2022.09.12.506413

**Authors:** Brendan O’Fallon, Ashini Bolia, Jacob Durtschi, Luobin Yang, Eric Frederickson, Katherine Noble, Joshua Coleman, Hunter Best

## Abstract

Detection of germline variants in next-generation sequencing data is an essential component of modern genomics analysis. Variant detection tools typically rely on statistical algorithms such as de Bruijn graphs, Hidden Markov Models and regression models, often coupled with heuristic techniques and thresholds to identify variants. Here we describe a new approach that replaces these handcrafted statistical methods with a single, end-to-end deep learning model that directly infers germline haplotypes from short read pileups. Our model, called Jovian, frames variant detection as a sequence-to-sequence modeling task, akin to language translation, and employs a transformer-based architecture to translate alignment columns into two predicted haplotype sequences. After training with 17 whole genome sequences from Genome-in-a-Bottle cell lines, we demonstrate that this method learns to realign complex and ambiguous read mappings to produce accurate haplotype predictions, predicts variant genotypes and phase accurately, and leverages the local read context to inform predictions about a given position. We also demonstrate that a 2-dimensional positional encoding significantly improved precision of the detected variants. Compared to other callers, sensitivity and precision is higher than GATK HaplotypeCaller, but lower than DeepVariant and Strelka2.

## 1 Introduction

Accurate inference of the DNA sequence variants in a sample is a fundamental element of genomics analysis. Tools that aim to identify variants must overcome several challenges to produce an accurate set of variant calls. For instance, the sequenced reads themselves may contain errors in the form of miscalled bases or PCR-induced repeat length errors, or the tools used to align reads to the reference genome may produce incorrect mappings (e.g. Li et al. 2014, Goldfeder et al. 2016). In addition, large or complex sequence variants may be obscured by limitations in the read alignment algorithm, and require a sophisticated assembly of reads to identify. Reconstructing real variation and discriminating it from technical error is challenging, and modern variant detection tools employ a variety of statistical and heuristic techniques to achieve high precision and sensitivity (e.g. De Pristo et al. 2011, Nielson et al. 2011).

All variant discovery tools must address two challenges. First, callers must identify the potential alleles in a region. Then, given a set of potential alleles, they must assess the likelihood of each candidate (or pairs of candidates in the diploid case), to determine which should be included in the caller output. Early callers, such as samtools / mpileup (Li et al. 2009) and the UnifiedGenotyper tool from the Genome Analysis ToolKit (GATK, De Pristo et al. 2011) rely on the read aligner to generate candidate alleles, and utilize several ad-hoc heuristics and thresholds to determine which alleles are most likely (e.g. Nielsen et al. 2011). Later tools (e.g. Poplin et al. 2018, Kim et al. 2018, Cooke et al. 2021) incorporated local re-assembly of reads in each region of interest, resulting in a significant improvement in the sensitivity and precision of variant calls. For instance, the HaplotypeCaller tool identifies candidate haplotypes by constructing de Bruijn graphs from k-mers present in the reads, and assesses haplotype likelihoods with a pair Hidden Markov Model (HMM).

More recent tools have incorporated elements of deep learning into the allele likelihood calculation or allele generation steps. DeepVariant (Poplin et al. 2018) uses a statistical method based on the HaplotypeCaller approach to identify candidates, but adds a Convolutional Neural Net (CNN) to classify variants as true or false positive detections. HELLO (Ramachandran et al. 2020), designed to work on hybrid short- and long-read datasets, employs a mixture-of-experts approach with separate 1-dimensional convolutions across the read and position dimensions of the input. Clair (Luo et al. 2019, Luo et al. 2020), uses a novel multitask deep learning approach to predict several properties of a potential variant at a given site, including zygosity and allele length.

Here, we describe a new approach that replaces both the candidate allele generation and the likelihood calculation with a single, learned model. We recast variant detection as a sequence-to-sequence translation problem in which the input sequence is the series of bases aligning to a single genomic location, and the output sequences are the two predicted haplotypes. The sequence-to-sequence translation is learned by a transformer-based deep neural network (Vaswani et al. 2017). This approach does not require any of the statistical machinery and heuristic thresholds utilized by most modern callers, and instead uses a deep learning model to reconstruct complete haplotypes directly from aligned NGS reads.

## 2 Methods

### 2.1 Model architecture & read encoding

Our model consists of a transformer-based encoder and a simple, fixed-length decoder that produces two output sequences, one for each predicted haplotype. The encoder component is an unmodified transformer with GeLU activations as implemented in PyTorch 1.10 (Vaswani et al. 2017, Paszke et al. 2019). Input tokens are the collection of bases that align to a given reference position, with some modifications described below. We add an additional fully connected layer prior to the transformer encoders which embeds the encoded basecalls in *d* dimensions, where *d* = 12 for the analyses here. The embedded basecalls are ‘flattened’ along the read dimension, producing an input token with size *dr* where *r* is the number of reads (*r* = 100 for all experiments proposed here). Instead of the typical 1-dimensional positional encoding, we employ a 2-dimensional encoding, allowing the model to differentiate across both the token (position) and read dimensions. The 2D encoding implementation follows Wang & Jyh-Charn (2021).

The decoder consists of a single fully-connected layer with ReLU activation followed by two additional fully-connected components that each generate a single haplotype. The decoder is not autoregressive and does not utilize special start-of-sequence or end-of-sequence tokens. Predictions are discrete probability distributions over the four possible bases at each predicted output position, produced with a softmax function over the raw outputs. The decoder is of fixed length and produces two haplotype sequences equal in length to the number of input tokens *g*. The full model architecture is shown in Figure 1.

**Figure 1:**
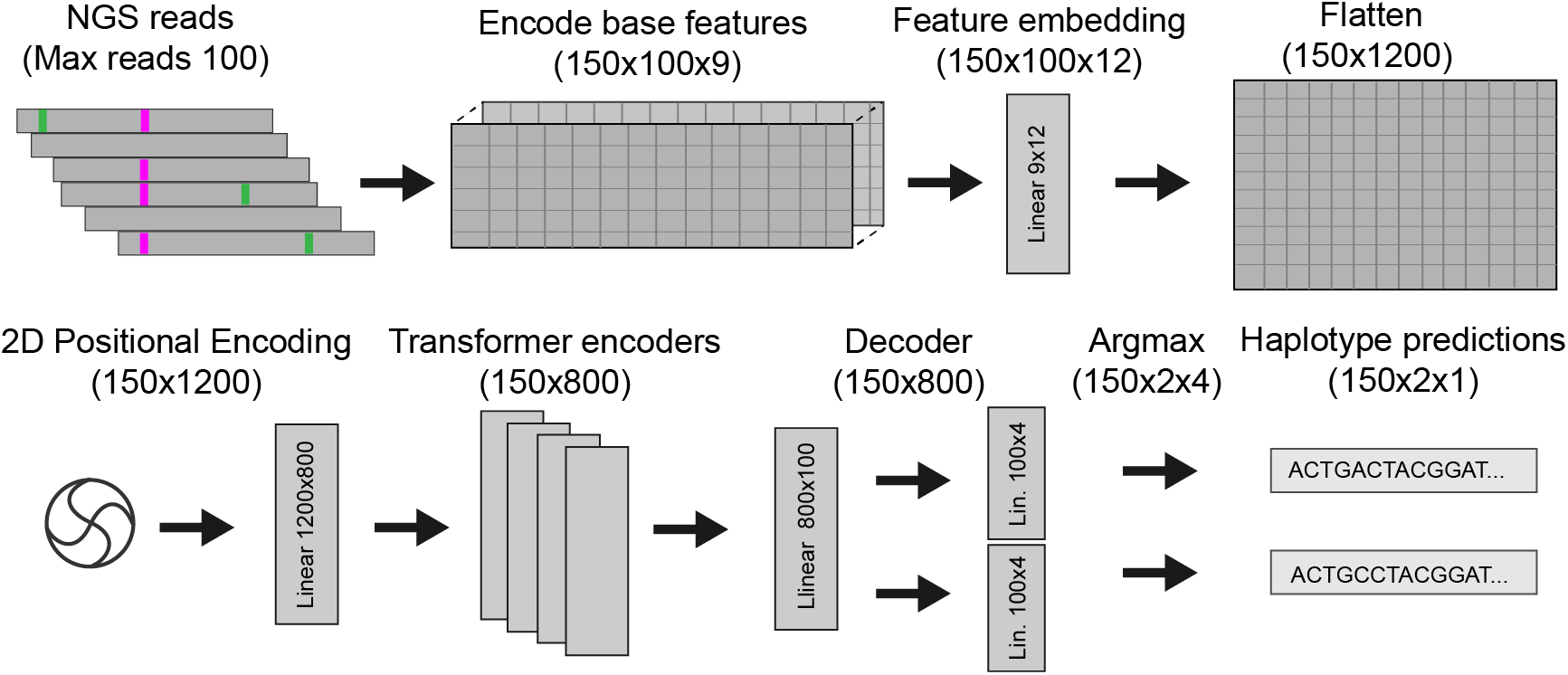
Overview of network architecture, numbers in parentheses indicate size of tensors set to zero.

Input tensors were generated by selecting a 150bp region from a set of aligned reads in BAM / CRAM format. For each region, all reads overlapping the window were obtained from the corresponding alignment file. Regions containing more than the maximum number of reads (100 for all analyses here) were downsampled to the maximum size. Reads were sorted by the reference coordinate of the first aligned base. For each aligned base in the selected reads nine features are encoded; the first four are the one-hot encoded base call, followed by base quality, two flags indicating if base ‘consumed’ reference base (i.e. was not an insertion) or consumed a sequence read base (was not a deletion), and additional flags indicating sequence read direction and clipping status. No gap tokens or other special handling was performed for insertions or deletions. In addition to sequenced reads, the first row in each encoded region was the reference sequence. For this special row we inserted a base quality of 100 for every position and did not set any of the other flags. Resulting tensors had dimension [*g, r*, 9], where *g* indexes genomic positions, *r* indexes reads, and 9 is the number of features. Positions in the input not corresponding to an aligned base were all

### 2.2 NGS data

Training data was obtained from 17 whole genomes sequenced from 5 Genome-in-a-Bottle cell lines. Eight samples were prepared with Illumina Nextera DNA Flex kit and the remainder were prepared with the Illumina TruSeq PCR-free kit. All samples were sequenced on an Illumina NovaSeq 6000 instrument in 2×150 mode, to an approximate read depth of 50. After conversion to fastq, the sequenced reads were aligned to human reference genome GRCh37 with the GEM-mapper (v3, Marco-Sola et al. 2012) and were sorted and converted to CRAM format with samtools version 1.9 (Li et al. 2009). No additional refinements, such as duplicate read marking, base-quality score recalibration, or indel realignment were performed.

### 2.3 Training data

To select regions to include for training data, we developed a scheme to sample regions in a biased manner, prioritizing regions containing variants and, especially, regions with multiple or complex variants. Regions of the reference genome overlapping the high-confidence regions from Genome-in-a-Bottle were subdivided into 150bp nonoverlapping windows, and these regions labelled according to the presence of variants. Separate labels were generated for regions containing a single SNV, deletion, or insertion, as well as regions containing multiple insertion-deletions, or those containing variants intersecting low-complexity regions. We additionally included ‘true negative’ regions where no known variant was present. In regions containing more than 100 overlapping reads, the reads were repeatedly downsampled to generate multiple training regions. For all analyses described here, we obtain training data only from the human autosomes 1-20, and hold out chromosomes 21 and 22 for model evaluation. Approximately 1.3M regions were obtained from each of the 17 samples, generating a full training data size of 24.3M regions. In addition to the “Full” training set we also examine a half-sized training data set of 12.1M regions, and quarter-sized set of 6.0M regions. The number of regions generated for each variant type is shown in Table 1.

**Table 1:** Number of regions selected from each sample to generate training data

| **Region type** | **Number of regions (per sample)** |
| --- | --- |
| No variant | 200,000 |
| SNV | 200,000 |
| Insertions | 200,000 |
| Deletions | 200,000 |
| Insertion/Deletion + SNV | 50,000 |
| Insertion + Deletion | 30,000 |
| SNV (low complexity) | 80,000 |
| Insertion (low complexity) | 80,000 |
| Deletion (low complexity) | 80,000 |
| Insertion + Deletion (low complexity) | 20,000 |
| Insertion/Deletion + SNV (low complexity) | 80,000 |

Target haplotype sequences were produced by obtaining truth variants from the Genomein-a-Bottle VCF files for each sample and inserting the variants into the reference sequence. Two sequences were generated for each region, one representing each haplotype. In regions where phasing of the variants was ambiguous, the reads in the sequenced sample were examined to determine phase status. Briefly, all possible genotypes (pairs of haplotypes) were generated and reads were aligned via Smith-Waterman to the possible haplotypes, and the highest scoring genotypes selected as the most likely phasing. This phasing procedure was only attempted for variants less than 100bp apart, otherwise the region was discarded.

### 2.4 Training procedure & loss function

Models were trained for 50 epochs unless noted otherwise, using the Adam optimizer, a learning rate of 5*e*^*−*5^, and batch size of 512. Models were implemented in PyTorch (v1.10, Paszke et al 2019).

For a given haplotype we use a modified cross-entropy loss function to compare the predictions to the target sequence. For each region two haplotypes exist and they have no inherent order. For instance, a single heterozygous variant can appear on either haplotype, and the model cannot learn to predict which because the assigment is random. To make the predictions opaque to haplotype order, we compute the loss in both haplotype configurations - predicted haplotype 0 with target haplotype 0 and 1 with 1, as well as predicted haplotype 1 with target haplotype 0, and 0 with 1, and backpropogate gradients only for the configuration with the lowest loss.

### 2.5 Variant detection

Given an alignment file in BAM or CRAM format, we first identify regions where a potential variant might exist. Any genomic position in which at least three reads contain a base that differs from the reference or an indel are flagged as potentially containing a variant. Positions closer than 100bp are merged into a single region.

For a single region containing suspected variants, we perform multiple overlapping forward passes of the model with step size *k*, where *k* = 50 for the results reported here. On each forward pass the model predicts two haplotypes by taking the base with the highest predicted probability at each position. Each haplotype is aligned via Smith-Waterman to the reference sequence, and any mismatching positions are converted to variant calls. After variants are collected across multiple windows, results are merged using a method that attempts to minimize the number of conflicting calls between windows. For each variant we record the number of windows in which each variant was detected, the number of variants in cis and trans, the mean probability of the variant bases, and the position of the variant within each window.

### 2.6 Variant quality calculation

The above variant detection procedure has two shortcomings. First, if every potential variant is emitted, then precision is poor. Second, the model does not innately produce well calibrated variant quality scores. To address these shortcomings we introduce a post-hoc random forest classifier that we train to discriminate true and false-positive variant calls (see table 2.6 for list of features). The resulting score is used as a final variant quality score suitable for tuning the tradeoff between sensitivity and specificity.

**Table 2:**
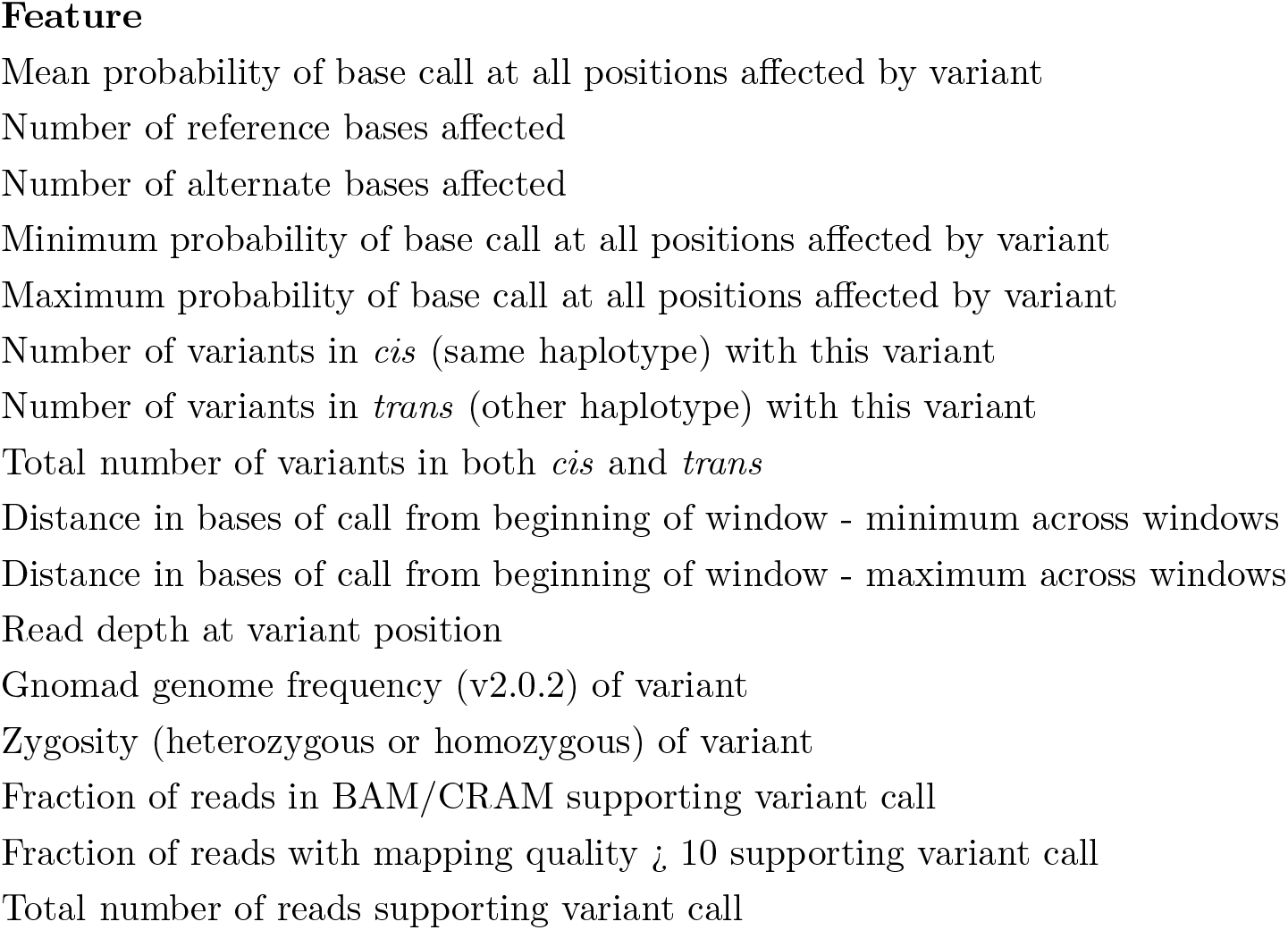
Features used to train classifier model for estimation of variant quality scores

The random forest classifier is trained by calling variants on selected regions from autosomes 1-4 totalling approximately 120Mb. True and false positive variant calls were identified with the *vcfeval* tool (Cleary et al. 2015). We used the scikit-learn implementation of the random forest with 100 trees, a maximum tree depth of 25, and ‘balanced’ class weighting option. Features are listed in Table 2.6.

## 3 Results

### 3.1 Model size and training data experiments

We explore the effects of model size and training data size on variant detection accuracy. Throughout, we calculate sensitivity and precision on a small set of evaluation regions from chromosomes 21 and 22 that were not used for training. Importantly, for performance reasons these sensitivity and precision calculations do *not* include the full variant detection procedure with multiple overlapping windows and the post-calling classifier, and instead reflect results from a single inference procedure on the target window (accuracy of the full detection procedure is explored below). We evaluate three model sizes and three sets of training data (Full, Half, and Quarter-sized, see section Training Data above for details).

During training, sensitivity and precision increased for all variant classes, with returns dwindling after 30-40 epochs of training (Figure 2). Little difference was observed between the Full and Half sized training data sets, but both sensitivity and precision were lower for the Quarter-sized training set. These results indicate that additional training examples beyond the Half size (12*e*^6^ example regions) offer little improvement in accuracy, and suggest that performance becomes limited by other factors, such as model capacity, inaccuracies in the training data set, or a non-optimal loss function.

**Figure 2:**
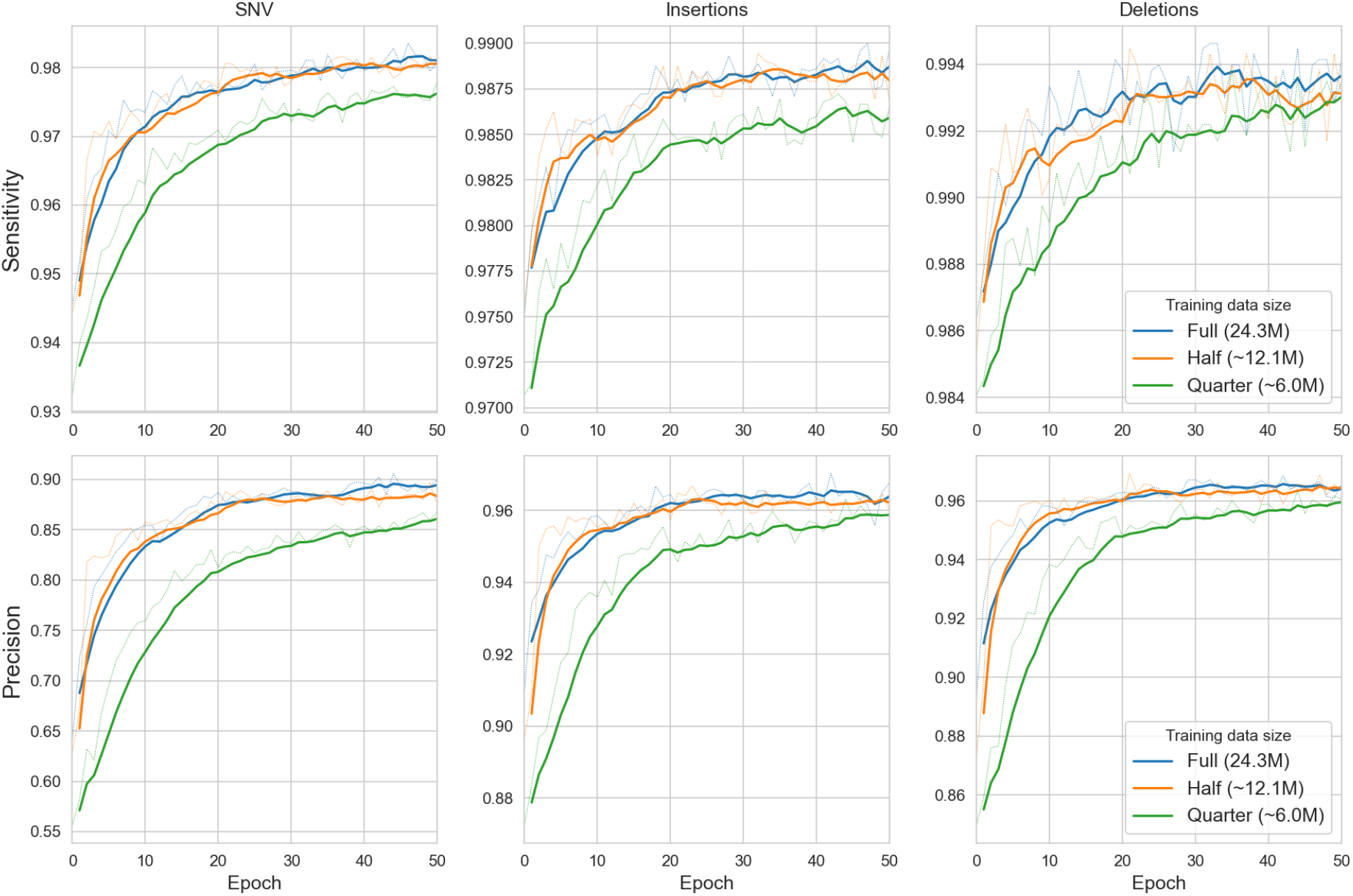
Sensitivity and precision curves for the 10M model during training for different training data sizes. Dashed lines are raw values, solid lines indicate rolling average with window size of 5.

Sensitivity was similar across all model sizes evaluated for all variant classes, but precision increased as model size was increased from 5M to 10M (Figure 3). However, precision did not increase further as model size was increased from 10M to 25M. Together with the training data size analysis, the results indicate that neither increasing model capacity beyond 10M parameters or training data size beyond approximately 12M examples results in improved model performance.

**Figure 3:**
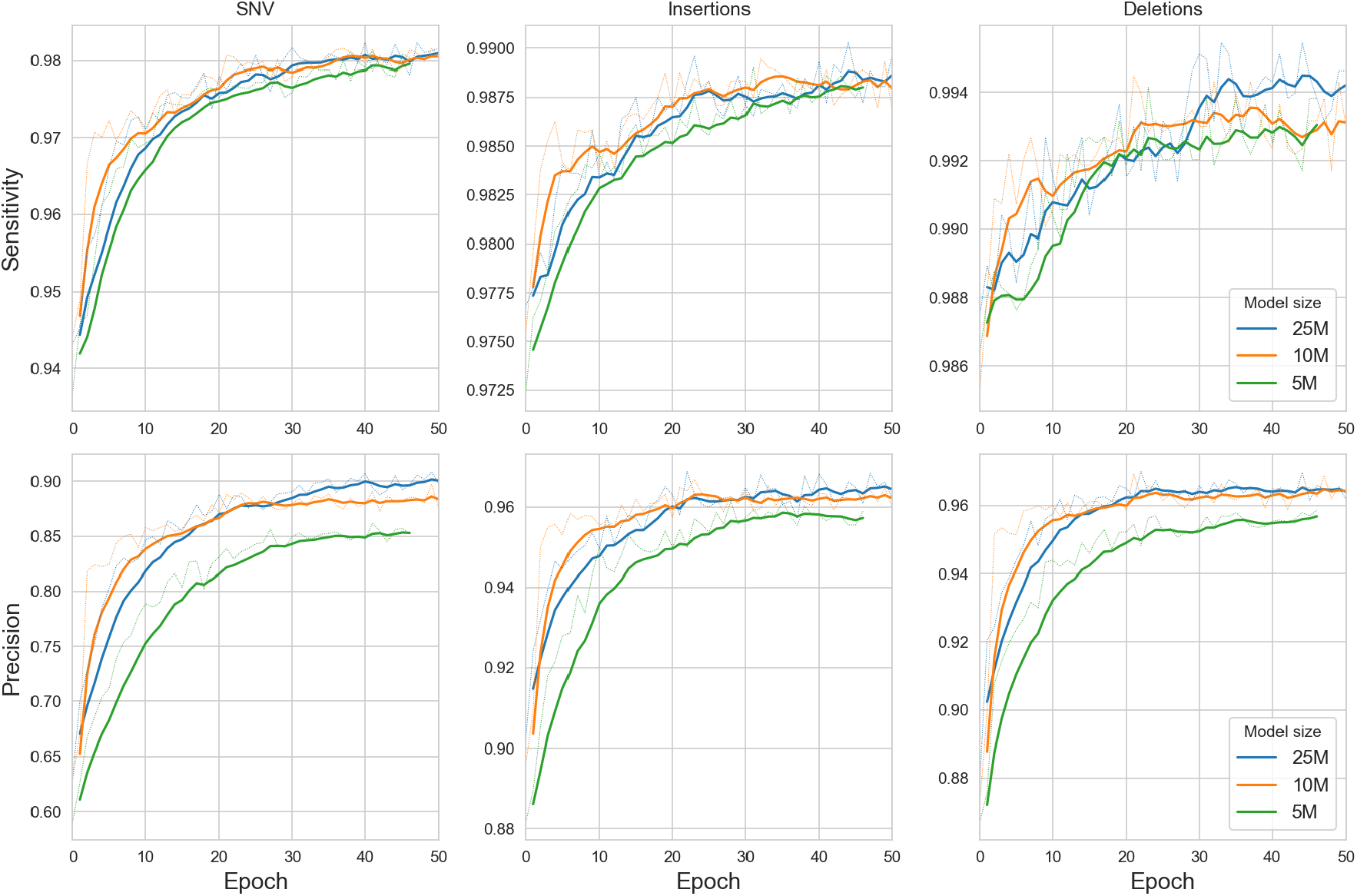
Sensitivity and precision curves for the Half-sized data set, for the 25M, 10M, and 5M model sizes. Dashed lines are raw values, solid lines indicate rolling average with window size of 5.

### 3.2 Variant detection accuracy

To characterize variant detection accuracy, we called variants on validation regions from chromosomes 21 and 22 for each of the 17 samples, and compared results to DeepVariant (v1.3.0, Poplin et al. 2018), HaplotypeCaller (v. 4.1.4.1), and Strelka2 (v. 2.9.10, Kim et al. 2018). For this comparison we used the 10M model trained on the Full set. We selected filtering criteria for each caller by reviewing ROC curves produced by *vcfeval* and selecting values close to the *F*_1_ maximizing value across samples. Jovian calls were filtered at quality 0.25, HaplotypeCaller at 60 (phred-scaled), Strelka at 4 (phred-scaled), and DeepVariant at 0. Sensitivity, precision, and related calculations were computed with the *hap*.*py* tool (Krushe et al. 2019) using Genome-in-a-Bottle (v4.2.1) as the benchmark variant set.

Across all regions, Jovian’s sensitivity and precision for indels is slightly higher than that for HaplotypeCaller, and slightly lower than DeepVariant or Strelka2 (Figure 4). For SNV calls Jovian has the highest sensitivity (99.1%) and precision similar to HaplotypeCaller (99.5%), but precision is lower than DeepVariant and Strelka2 (99.8% and 99.9%, respectively).

**Figure 4:**
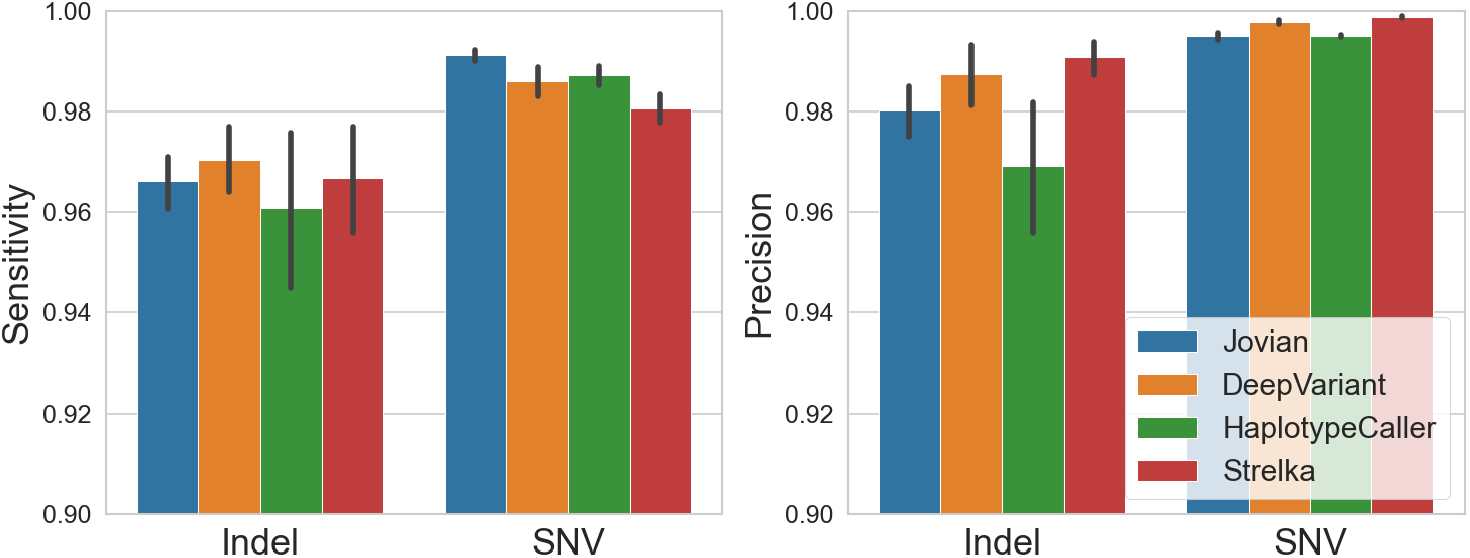
Sensitivity and precision of Jovian variant calls (10M parameter model, Full training size) compared to other callers

**Figure 5:**
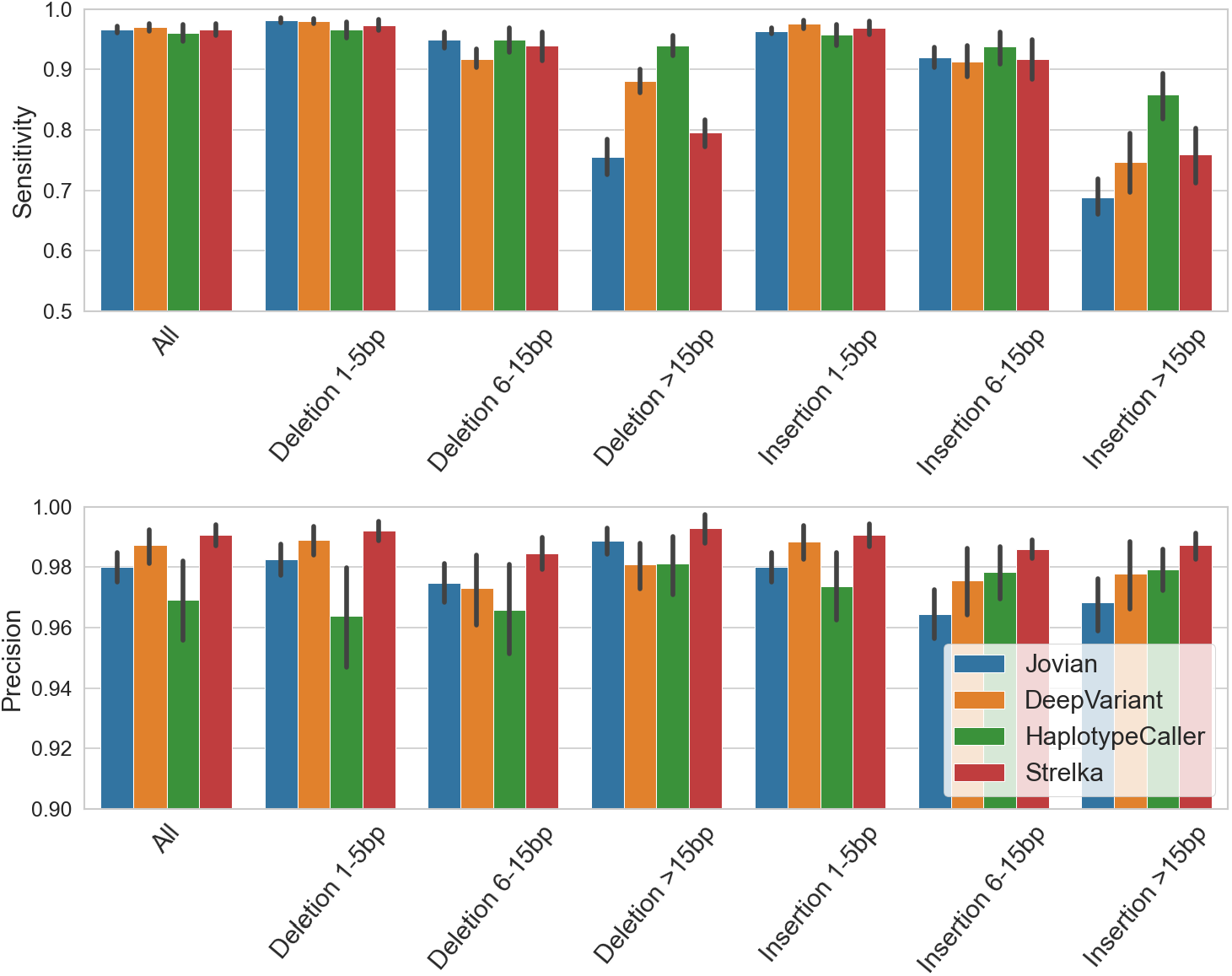
Sensitivity and precision of Jovian indel calls by size class (10M parameter model, Full training size) compared to other callers

Across indel size classes Jovian has high sensitivity for deletions less than about 15bp, but relatively low sensitivity for larger deletions (*≈*75%). For insertions Jovian has sensitivity similar to other callers at smaller sizes, but again reduced sensitivity for the largest class (*≈*68%). Precision follows a similar pattern, but with relatively high precision for smaller indel sizes and reduced precision at larger sizes.

## 3.3 Genotyping & Phasing

To check the accuracy of predicted genotypes and phasing of variants we compared calls in the chromosome 21 and 22 evaluation regions to the known genotype and phase from Genome-in-a-Bottle (GIAB) variant data. Approximately 99.5% of homozygous and 99.0% of heterozygous calls matched the true genotype. For phasing accuracy, we identified pairs of heterozygous variants less than 100 base pairs apart and compared their phasing predictions with the known phasing information from GIAB. The phasing information in GIAB samples is only available for HG001, HG002 and HG005 cell lines, so analysis was restricted to those samples. Phase predictions matched GIAB phase for approximately 96.9% of SNV pairs and 99.3% of indel pairs. Together, these analyses indicate that the model learns to reconstruct variant phase and genotype accurately, producing pairs of haplotype sequences that match those present in the sample with high fidelity.

### 3.4 Complex regions

While many genomic variants are trivial to identify by examination of mapped reads, some regions require a more sophisticated reassembly to identify the true haplotypes. We examine the ability of our model to perform this more complicated reconstruction by identifying real variants in regions containing multiple conflicting read mappings or extensive read softclipping.

To identify variants requiring complex reconstruction, we first obtained all indel variants from Genome-in-a-Bottle greater than 4 bp in length from the validation regions on chromosomes 21 and 22. For each variant, we examine the reads in the alignment file that overlap the variant position, and flag any with more than 4 distinct indel start sites or at least 20% reads with more than 10bp soft-clipped as requiring ‘complex’ reconstruction. These variants were moved to a separate truth set. After running our variant calling procedure on the alignment file, we computed the fraction of variants from the truth set that were identified by our model. This procedure was repeated for all 17 samples in our data set.

Our model was able to reconstruct the correct haplotypes for *≈*89.5% (mid 50% 82.5-94.8) of these variants, somewhat lower than other callers (table 3.4). These results indicate that the model learns a reconstruction procedure more nuanced than a simple column-by-column examination of reads. Instead, the model learns to synthesize information across tokens to produce complete haplotypes with accuracy similar to (but slightly less than) the top performing current methods.

**Table 3:** Model parameter combinations used for the three model sizes investigated

| Model name | Encoder layers | Attn. heads | Hidden dim. | Embedding dim. |
| --- | --- | --- | --- | --- |
| 5m | 4 | 4 | 200 | 600 |
| 10m | 6 | 6 | 250 | 400 |
| 25m | 8 | 8 | 400 | 800 |

**Table 4:** Sensitivity of detection algorithms for variants requiring complex reconstruction

| Caller | Sensitivity | Mid 50% |
| --- | --- | --- |
| Jovian(10M) | 89.5% | 82.5 - 94.8 |
| HaplotypeCaller | 95.8% | 91.0 - 100.0 |
| DeepVariant | 91.7% | 83.5 - 97.1 |
| Strelka | 93.1% | 90.9 - 98.4 |

### 3.5 Use of local context for SNV prediction

Variant callers must integrate multiple types of information to produce accurate calls, including the fraction of reads supporting a given call (Variant Allele Frequency, VAF),base and mapping qualities, read position, and potentially information from nearby positions. To examine the influence of the number of nearby reference mismatches, VAF, and base quality on the likelihood that a prospective SNV was called, we first collected genomic positions in which 10 - 30% of reads supported a non-reference basecall. We then tallied the number of genomic positions at which 10% - 20% of reads supported a non-reference call in a window 20bp upstream and downstream of the candidate. We also stratified the candidate position into low base quality and high base quality groups, and smaller VAF windows of 10-20%, 15-25% and 20-30%.

The probability that a prospective variant is called is strongly associated with the number of nearby alignment columns containing mismatches (Figure 6). For instance, prospective variants with VAF between 20-30% and high base quality were called in over 60% of examined cases when there were no nearby mismatches, but called in less than 30% of cases when a single other column contains a mismatch, and in less than 10% of cases when 5 or more nearby positions contain a mismatch. A similar, but less pronounced trend exists when the prospective variant is at lower VAF (left two panels, figure 6). When the prospective variant has poor base quality scores, the association disappears and the likelihood that the variant is called remains uniformly low. Overall, these results indicate that the model learns a nuanced SNV detection procedure that discards most prospective SNVs with low base quality, but selectively retains those with higher base quality provided that not too many nearby positions contain reference mismatches.

**Figure 6:**
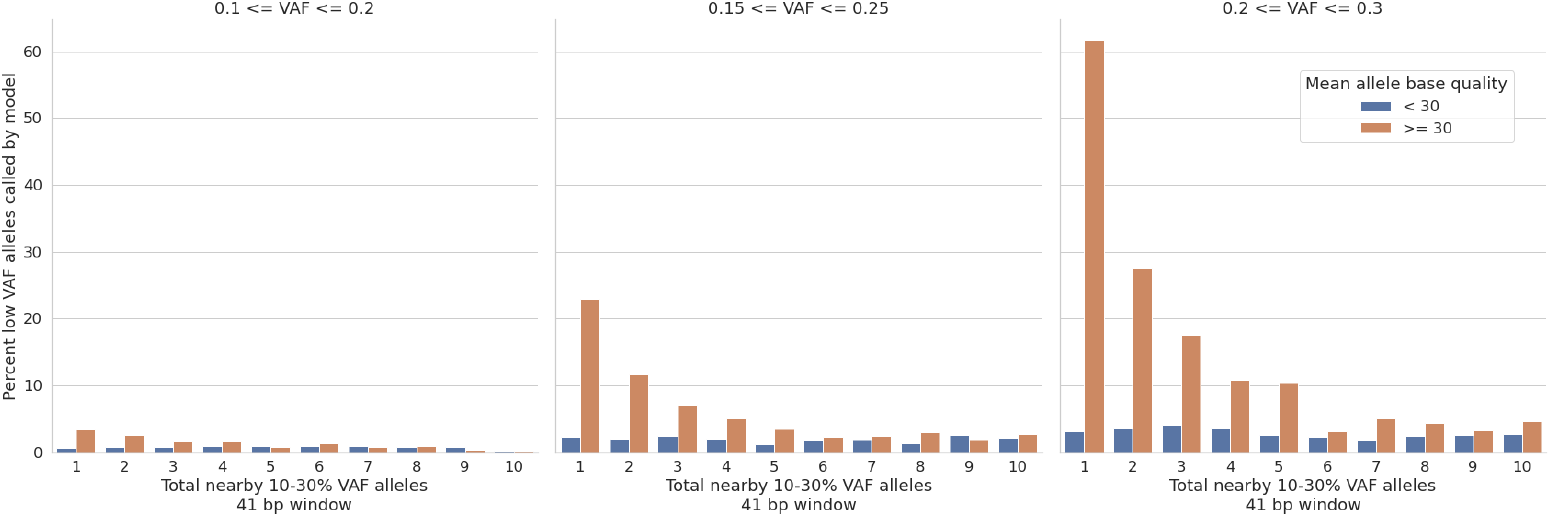
Effect of number of nearby reference mismatches on the likelihood that a poorly supported candidate variant will be called.

### 3.6 Ablation of 2D positional encoding

To assess the impact of the two-dimensional positional encoding, we replaced the 2D encoding module with traditional 1D encoding, and left other model parameters unmodified. We conducted two training runs, one with the 10M model and Quarter training data size, and the other with the 25M model and Half training data size. Both training runs were conducted for 25 epochs. When compared to their equivalent models with 2D encoding, the 1D encoding model showed substantially reduced precision, in particular for the SNV class (Figure 7). Sensitivity was also reduced but to a lesser degree.

**Figure 7:**
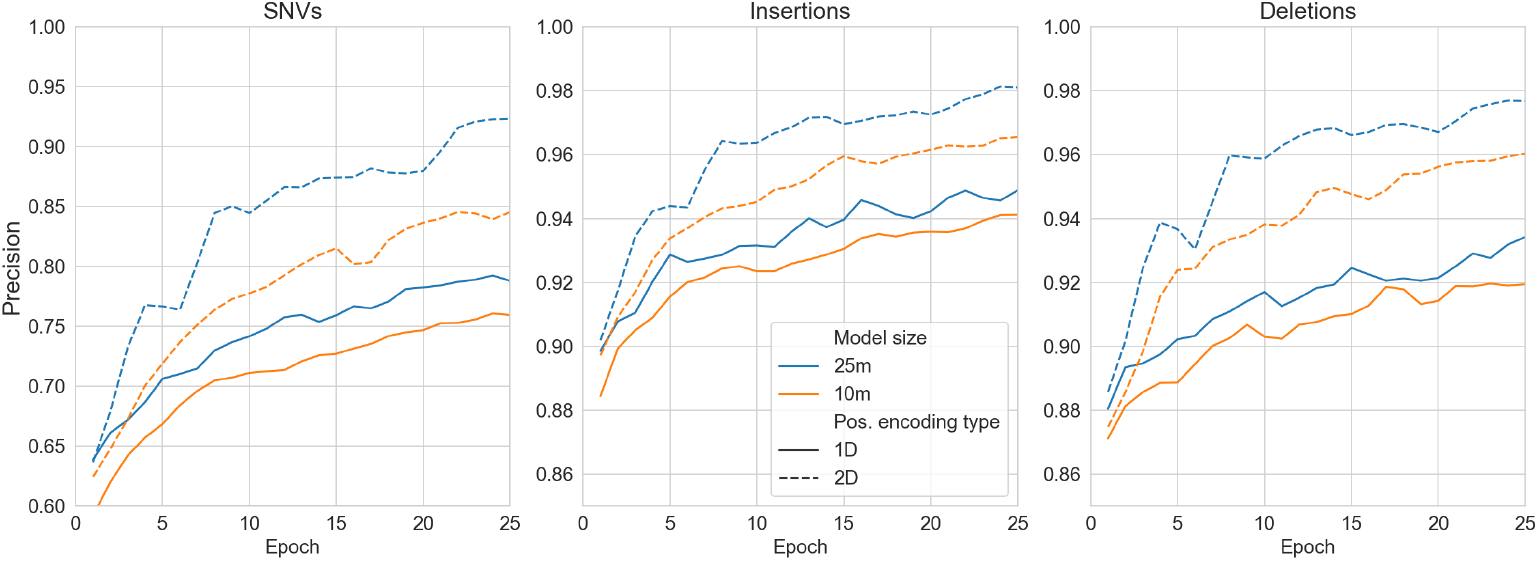
Precision of variant calls for each variant class for 1-dimensional and 2-dimensional positional encodings, for two different model sizes

By applying positional encoding across both the read and position (input token) dimensions, two-dimensional encoding allows the model to track which read contains a given base across positions. We hypothesize that read-level tracking is important for precision because mismapped or poorly aligned reads generate many reference mismatches, most of which are likely to be errors. The model must learn to identify and ignore such reads, and 2D encoding facilitates this because the model can more easily track read-level features across input tokens.

## 4 Discussion

We describe a new approach to the problem of detecting sequence variants in next-generation sequencing (NGS) data. Our approach envisions variant detection as a sequence-to-sequence modeling problem, akin to language translation, and leverages the successful transformer architecture to perform the translation. The sequence-to-sequence approach allows a single deep learning model to be used for candidate allele generation and haplotype construction. In contrast to other variant detection methods (e.g. Poplin et al. 2018, Kim et al. 2018), our approach does not rely on any handcrafted statistical techniques such as de Bruijn graphs, hidden Markov models, or heuristic thresholding and instead learns to construct accurate haplotypes directly from aligned NGS reads.

Our analysis demonstrates that the model learns many features of modern, state of the art variant callers. For instance, our model learns to reconstruct large, complex variants in the presence of ambiguous mapping and extensive soft clipping with a sensitivity near 90%. In addition, the model learns to employ local context and read-specific features to predict phase and genotype accurately, and pairs of variants are phased correctly in 97-99% of cases where variants are within 100bp. The model also learns to consider the surrounding read context when predicting the presence of a SNV. For instance, a potential SNV in a region with few reads containing reference mismatches is much more likely to be called than a similar variant in a region where many reads contain mismatches, indicating that the model has learned to be cautious of potential SNVs in ‘dirty’ regions where many reads contain errors or are misaligned.

Overall variant detection accuracy is similar to other modern callers. At our chosen quality cutoffs (selected to be near the *F*_1_-maximizing value for each caller), Jovian’s performance on insertion-deletion variants outperforms HaplotypeCaller, but falls short of DeepVariant and Strelka2 (Figure 4). The differences are partially due to Jovian’s relatively poor sensitivity for indels larger than 15bp (mean 75.6% for deletions, 68.8% for insertions) and low sensitivity in tandem repeat regions (mean sensitivity 84.0%). For SNVs Jovian demonstrated the highest sensitivity among all callers (mean 99.1%), and precision similar to HaplotypeCaller but lower than DeepVariant and Strelka2 (Jovian mean precision 99.5%). To our knowledge, the transformer architecture has not been previously explored for variant detection, but a model similar to ours was proposed by Baid et al. (2021) for generation of accurate consensus reads from PacBio data. The Baid model (‘DeepConsensus’) used a network architecture similar to ours, but included an extra token representing a gap between the aligned sequences, and a novel loss function that accounted for gaps between the predicted and true sequence in a region. In contrast we use a simple cross-entropy loss function, do not introduce gaps in the aligned sequences compared to the reference, and use 2-D positional encoding.

In this work we have used a standard cross-entropy loss function that simply compares each prediction at base *i* to the label at base *i* for each haplotype. While training with this loss yields accurate results, it has two shortcomings. First, the impact of an insertion / deletion (indel) error depends on its position in the region, with indels at lower indices leading to a greater number of mismatches and high loss compared to indels at higher indices. Secondly, indel errors are punished more severely than similar SNV errors. We hypothesize the second fact contributes to the lower performance of our model on SNV calls compared to indels. Alternative, alignment-based loss functions such as those described by Baid et al. (2021) and Petti et al. (2021) may provide a path forward, however the Petti implementation lead to lower sensitivity and much longer training times in our experiments.

Our model uses an unmodified transformer encoder with a very simple decoder consisting of just two fully connected layers, and we leave to future work exploration of the many transformer variations proposed in recent years (e.g. Dosovitskiy 2018, Fedus et al. 2021, Liu et al. 2021, Wu et al. 2021). Our simple decoder has a fixed length equal to the number of input tokens (genomic positions), but a natural improvement would be to add a more sophisticated decoder capable of producing a variable length sequence to handle larger insertion and deletion events. In addition, newer transformer models that avoid the *O*(*n*^2^) scaling with the number of input tokens may be able to handle longer input sequence lengths, and therefore may be able to detect larger variants.

## 5 Availability

Source code for is available via git at https://github.com/brendanofallon/jovian

